# Pharmacophore modeling, 2D-QSAR, Molecular Docking and ADME studies for the discovery of inhibitors of PBP2a in MRSA

**DOI:** 10.1101/2025.02.13.636556

**Authors:** Fredrick Mutie Musila, Grace Wairimu Gitau, Peris Wanza Amwayi, James Munyao Kingoo, Dickson Bennet Kinyanyi, Patrisio Njiru Njeru

## Abstract

Methicillin-resistant *Staphylococcus aureus* (MRSA) is considered to be a worldwide threat to human health and the global spread of MRSA has been associated with the emergence of different types of infections and resultant selection pressure due to exposure to many antibiotics. In the current era characterized by incessant antibiotic resistance, assessment of multiple molecular targets represents notable therapeutic opportunities in the medical and pharmaceutical industry and can aid in the discovery of novel molecules that inhibit various receptors effectively to replace the current weak antimicrobial agents. Penicillin binding protein 2a (PBP2a) of MRSA is a major determinant of resistance to β-lactam antibiotics. The activity of PBP2a is not inhibited by β-lactam antibiotics, allowing the strain to survive in the presence of β-lactams leading to resistance to β-lactam antibiotics. The study aimed at identifying potential inhibitors of PBP2a receptor of MRSA through ligand-based pharmacophore modeling, 2D-QSAR, structure-based drug design docking and molecular dynamic (MD) simulations. The study led to the development of a satisfactory, predictive and significant 2D-QSAR model for predicting anti-MRSA activity of compounds and also led to the identification of two molecules: C_21_H_25_N_7_O_4_S_2_ (ChEMBL30602) and C_20_H_17_NO_6_S (ChEMBL304837) with favorable pharmacophore features and ADME properties with potential to bind strongly to PBP2a receptor of MRSA. MD simulation analysis showed that the interactions of C_20_H_17_NO_6_S (ChEMBL304837) with PBP2a over 100 ns was more stable and similar to the interaction of ceftobiprole with PBP2a and may become potential drug candidate against MRSA which has developed a lot of resistance to current antibiotics.

## Introduction

Certain variants of *Staphylococcus aureus* have developed resistance to methicillin, a penicillin derivative altering methicillin-susceptible *staphylococcus aureus* (MSSA) to methicillin-resistant *staphylococcus aureus* (MRSA). The emergence of MRSA was first identified in 1960 in patients following the administration of methicillin to treat penicillin-resistant *Staphylococcus aureus* strains (Turner et al., 2019). The spread of MRSA poses a serious challenge in the treatment of severe infections and according to recent World Health Organization reports in 2017, MRSA is considered to be a worldwide threat to human health (Frieri et al., 2017). Subsequently, the global spread of MRSA has been associated with the emergence of different types of infections and the resultant selection pressure exerted by exposure to various antibiotics (Harkins et al., 2017).

Various strains of MRSA have been reported such as hospital-associated MRSA (HA-MRSA) clonal complex 30 (CC30) in North America, the United Kingdom and Europe, community-associated MRSA (CA-MRSA) USA300 in North America, livestock-associated MRSA (including ST398 and ST93) in Australia (Casey et al., 2013; David et al., 2014; Kennedy et al., 2008; McAdam et al., 2012; Turner et al., 2019) and infections in wild animals such as Hedgehog in Denmark (Larsen et al., 2022).The HA-MRSA and CA-MRSA infections differ in terms of their clinical manifestation, molecular biology, antibiotic susceptibility and response to treatment (Siddiqui & Koirala, 2021). HA-MRSA infections are one of the major causes of healthcare-acquired infections worldwide and are often associated with prolonged hospitalization that leads to increased health cost burden, significant high morbidity and mortality rate (Elward et al., 2009; Lakhundi & Zhang, 2018; Shahkarami et al., 2014). Furthermore, HA-MRSA infections are endemic in healthcare facilities and this is a serious impediment to its eradication (Grema et al., 2015).

MRSA is a consequence of the acquisition of a staphylococcal chromosomal cassette mec (SCCmec) mobile genetic element that is integrated into the *Staphylococcus aureus* chromosome. SCC*mec* comprises methicillin-resistant determinant *mecA* gene and a set of site-specific recombinase genes (*ccrAB* and *ccrC*) responsible for mobility (Turner et al., 2019). The *mecA* gene encodes the enzyme transpeptidases known as penicillin-binding protein 2a (PBP2a) which is also designated as PBP2′. PBP2a is a major determinant of resistance to β-lactam antibiotics. It plays an essential role in the crosslinking reaction of the peptidoglycans which is a principle building unit in the bacterial cell wall during biosynthesis of a mature cell wall and maintenance (Bugg & Walsh, 1992; Katayama et al., 2000; Meroueh et al., 2006). The activity of PBP2a is not inhibited by β-lactam antibiotics, hence permitting the strain to survive in the presence of β-lactams leading to resistance to β-lactam antibiotics (Fishovitz et al., 2014). On the contrary, other PBPs that perform the transpeptidase activity can be inhibited by β-lactam antibiotics. Additionally, mutation of PBP in *S. aureus* is suggested to contribute to methicillin resistance which is transferred across *S. aureus* organisms by bacteriophage transduction (Lakhundi & Zhang, 2018). Another mechanism of beta-lactam resistance in MRSA is through the synthesis of beta-lactamases encoded by the *blaZ* gene, which is responsible for the hydrolysis of the amide bond of the four-membered beta-lactam ring. The hydrolysis contributes to the inactivation of the beta-lactams before interaction with the PBP target (Fisher & Mobashery, 2020; Sandanayaka & Prashad, 2002)

PBP2c and blaZLGA251-encoded penicillinase are orthologues of the PBP2a enzyme and show amino acid sequence identity of 63% and 65% respectively when aligned with PBP2a (García-Álvarez et al., 2011). Penicillinases appear to have a narrower spectrum that provides resistance only to penicillin G and other penicillinase-labile subclasses of penicillin compared to PBP2a and PBP2c enzymes (Larsen et al., 2022). Even though PBP2a is a well-established antibiotic target of MRSA, it has low binding affinity for transpeptidase-inhibiting β-lactam antibiotics. This reduced affinity prevents β-lactams from inhibiting MRSA cell wall synthesis therefore resulting in the continued multiplication of the bacteria in the presence of β-lactams leading to the of MRSA resistance (Fergestad et al., 2020). Therefore, there is a need for the development of compounds that can effectively inhibit PBP2a and circumvent MRSA antibiotic resistance.

Currently, vancomycin is the preferred antibiotic for the treatment of severe MRSA infections. Other alternative antibiotics such as linezolid and daptomycin with higher efficacy than vancomycin, are also used as substitute treatment options (Arbeit et al., 2004; Fowler Jr et al., 2006; Shorr et al., 2005; Weigelt et al., 2005). However, recent reports indicate the susceptibility of bacteria is limited by continuous recurrent septicemia, adverse effects on renal function, and high probability of emergence of bacteria resistance to these antibiotics (Aljohani et al., 2020; Holmes et al., 2012; Kshetry et al., 2016). MRSA consists of qacA and qacB genes that code for multidrug endogenous efflux protein Qac A and Qac B, respectively which are found in the cell membrane. These proteins have been linked with decreased susceptibility to non-β-lactam antibiotics and chlorhexidine antibiotics (Foster, 2016; Hong et al., 2019; Jang, 2016). Furthermore, HA-MRSA resistance to other categories of antibiotics like aminoglycosides, lincosamides, fluoroquinolones, and macrolides that are currently in clinical use has often been reported (Kot et al., 2020).

Decreased development and release of novel antibiotics as well as the prime focus of current antibiotic development pipelines on antibiotics combination therapy to decrease antibiotic resistance, pose a threat to global efforts to contain drug-resistant infections (Tyers & Wright, 2019). Therefore, the discovery of inhibitors of the well-characterized targets that lead to antibiotic resistance is vital (Lade & Kim, 2021). Two effective new cephalosporins that effectively bind to PBP2a of MRSA are ceftaroline and Ceftobiprole (Chan et al., 2015; Long et al., 2014). These two antibiotics are also able to bind to PBPs in gram-positive bacteria including those resistant to other β-lactam antibiotics. However prior studies have identified ceftobiprole and ceftaroline-resistant methicillin-resistant hence the need to search for more effective antibiotics (Chan et al., 2015).

*In silico* approaches provide a cost-effective way of identifying new potential antibacterial drugs. In this study, pharmacophore modeling, quantitative structure activity relationship (QSAR) and molecular docking were used to identify new potential ligands for the PBP2a receptor of MRSA that is responsible for resistance to beta-lactams. QSAR involves the development of a machine-learning model which can be used to predict the activity of unknown compounds. The model is developed using compounds whose activities are known and well characterized to determine the activity of the unknown compounds. Pharmacophore modeling involves looking for common features of several drugs known to block a particular receptor referred to as the pharmacophore of drugs. In this study, the pharmacophore features were utilized to search for other compounds across various databases that appear to potentially block the PBP2a as the receptor of interest. The activities of the compounds were then predicted using the QSAR model. Pharmacophore modeling, 2D-QSAR and virtual screening studies have previously reported the use of this approach toward the identification of novel inhibitory leads targeting glycogen synthase kinase 3b (Taha et al., 2008), influenza neuraminidase (Abu Hammad & Taha, 2009), β-secretase (A. Al-Nadaf et al., 2010) renin (A. H. Al-Nadaf & Taha, 2011), glycogen phosphorylase (Taha et al., 2011), β -glucosidase (Khalaf et al., 2011), and β-galactosidase (Abdula et al., 2011), cyclin-dependent kinase (Al-Sha’er & Taha, 2010) besides myriad of other studies. Since multi-drug resistance has become a major drawback in the management of MRSA infections, the aim of this study was the identification of potential inhibitors of PBP2a of methicillin-resistant *staphylococcus aureus* through ligand-based pharmacophore models, 2D-QSAR, ADME screening, structure-based drug design docking and MD simulations.

## Materials and Methods

### Selection of Data for QSAR

Data for QSAR was obtained from BindingBD (https://www.bindingdb.org/) site which is an online public database of measured binding affinities, centered mainly on the interactions of proteins considered to be candidate drug-targets with small drug-like molecules (Gilson et al., 2016). Using PBP2a as the target protein, a search of various molecules/ligands that are known to bind to PBP2a as the target protein was done. Data obtained from BindingBD included parameters such as equilibrium dissociation constant (Kd), inhibitor constant (Ki), half-maximal inhibitory concentration (IC_50_) and half-maximal effective concentration (EC_50_), of the molecules against the PBP2A.

### Generation of 2D descriptors

2D Descriptors were generated using PaDEL which is a tool used for calculation of molecular descriptors and fingerprints (Yap, 2011). Using the ligands in SMILES format, PaDEL was able to generate 2D descriptors which were then combined with the ligands’ IC_50_ values from the bindingBD. Descriptors acted as the independent variables while IC_50_ values acted as the dependent variable during 2D-QSAR model development.

### QSAR model development

The 2D-QSAR model was developed in R statistical program using Multiple Linear Regression (MLR). Data was first cleaned through the removal of descriptors with blank entries (blanks), those with identical values (constant columns) and as well as removing correlated columns. Data was split into training set and test set in the ratio of 6:4 respectively then the 2D-QSAR model was built using a forced entry approach under MLR. The model developed was used to predict the anti-PBP2a activity of structurally similar molecules thought to bind to the PBP2a receptor of MRSA.

### QSAR Model evaluation and validation

The developed model was internally and externally moderated using test and new data to ensure its stability, generalization and predictability (Gramatica et al., 2013). Model summary statistics which are needed for model valuation and validation were also investigated and included F statistic, Multiple R^2^, Adjusted Multiple R^2^, Durbin Watson Statistic, Variable Inflation Factor (VIF), Mean absolute error (MAE) Mean Bias Error (MBE), Relative Absolute Error (RAE), Mean Absolute Percentage Deviation (MAPD), Mean Squared Error (MSE) and Relative Mean Squared Error (RMSE) (Puzyn, 2022; Veerasamy et al., 2011).

### Pharmacophore modeling

Two drugs (ceftaroline and ceftobiprole) which are used to treat MRSA by specifically binding to PBP2a were obtained from Pubchem and used for the identification of pharmacophore features (Kim et al., 2016). Similarly, five ligands that had the highest activities against PBP2a of MRSA from BindingBD were also used. All seven ligands were analyzed in PharmaGist Webserver (https://bioinfo3d.cs.tau.ac.il/PharmaGist/) to identify the common best pharmacophore features for searching other molecules with similar features from publicly available databases (Schneidman-Duhovny et al., 2008).

### Molecule search and ADME Screening

Using the best pharmacophore features obtained from PharmaGist Webserver, other molecules with such pharmacophore features were searched using Pharmit (http://pharmit.csb.pitt.edu/) across various chemical databases such as Molport, Zinc database, ChEMBL, ChemDiv, ChemSpace, MCULE, MCULE ULTIMATE, NCI Open Chemical Repository and LabNetwork (Sunseri & Koes, 2016). Molecules obtained were then subjected to ADME screening which also included Pan-Assay Interference Compounds (PAINS) (Baell et al., 2013) and Brenk checks (Daina et al., 2017) to eliminate toxic compounds or those likely to give false positives during *in vitro* screening. Drug likeness, pharmacokinetics, lipophilicity and physicochemical parameters were also predicted. The final list of molecules with favorable ADME properties was then subjected to the 2D-QSAR model prior built for the prediction of their anti-PBP2A activity.

### Molecular docking

From the prediction based on the 2D-QSAR model, the best hits were selected and subjected to molecular docking. The 3D structure of PBP2a was obtained from Protein Data Bank (PDB ID: 5M18) and together with the selected ligands prepared in UCSF Chimera (Pettersen et al., 2004). Ligand preparation was done through effective and robust compilation of UCSF chimera to produce globally minimized conformers with profound properties important for drug-likeness. Generally undesirable water molecules and other heteroatoms are deleted, hydrogen atoms are added. Indeed, ligand preparation involves, penetration into the quantum chemistry and location of exact tautomeric isomers and fixing them non-canonically, neutralization of charged groups, generation of alternative chiralities and optimization of geometries. Preparation of the PBP2a receptor also involved the removal of water, heteroatoms and the co-crystalized ligand(s). Identification of the active sites of PBP2a was done in BIOVIA Discovery Visualizer (Biovia, 2017). Molecular docking was done in PyRx using AutoDock Vina and the number of docking runs was set at 9 for each ligand-receptor complex (Dallakyan & Olson, 2015). Biovia Discovery Visualizer was used to study the molecular docking results to identify various binding energies and binding interactions of the various ligand-receptor complexes.

### Molecular Dynamics Simulation

This study conducted a molecular dynamics (MD) simulation lasting 100 ns to stabilize the protein structure and also assess the binding efficacy of the compound to the target protein. The GROMACS 2022.4 package was used for conducting MD simulations on all complexes (Van Der Spoel et al., 2005). The topology of the molecules was determined using the CHARMM36 force field parameter (Jo et al., 2008), which was then applied to both the ligands and protein. The Cgenff server was employed to produce the topologies and parameter files for the ligand (Vanommeslaeghe et al., 2010). Furthermore, the Particle Mesh Ewald (PME) approach was employed for the calculation of electrostatic forces (Toukmaji et al., 2000). The solvation of each ligand-protein combination was performed using the transferable intermolecular potential with a three-point (TIP3P) water model. The complex was reconstructed using a cubic box, with a buffer distance of 1Å. The neutrality of the system was achieved through the incorporation of Na+ and Cl-ions into the system. The successful mitigation of unfavorable connections and collisions within the protein structure was achieved by implementing the energy reduction strategy and performing 5,000 iterations of the steepest descent method. The LINCS method (Hess et al., 1997) was employed to constrain the hydrogen bonds, after which the entire system was brought up to a temperature of 310 K. Following energy minimization, the complexes underwent two successive equilibration stages. The initial phase involved a 1ns equilibration under the NVT ensemble, followed by a further 1ns equilibration under the NPT ensemble. The velocity-rescaling approach was employed to incorporate temperature coupling (Bussi et al., 2009), while the Parrinello-Rahman pressure method (Martoňák et al., 2003) was employed to maintain constant pressure. During the 100 ns production run, a system that had reached equilibrium was employed. A comprehensive analysis of the complete system was conducted utilizing the several analytical modules provided by the GROMACS package, focusing on structural and conformational aspects. Moreover, post MD analysis included determination of RMSD, RMSF, SASA, Rg and the number of hydrogen bonds. Lastly, the binding free energy of the protein-ligand complex was determined using the GROMACS add-on tool gmx MM/PBSA (Paissoni et al., 2014). Here, the last 20 ns of the MD simulation was used for the calculation of the binding free energy.

## Results and Discussion

### 2D-QSAR model

From the BindingBD database, data from 31 compounds that are known to bind to PBP2a were extracted. The data included parameters such as kd, ki, IC50, and EC50 of the molecules against the PBP2a (**Supplementary_data_1.xlsx**). IC_50_ data which was used as the dependent variable for the 31 compounds together with 1446 descriptors (predictor/independent variables) generated from PADEL software (**Supplementary_data_2.xlsx**) were cleaned and used to develop 2D-QSAR model in R. After removal of descriptors with blank entries (blanks), identical values (constant columns) and as well as removing the correlated columns, only 16 descriptors were used to build the 2D-QSAR model. The 2D-QSAR model developed was evaluated and externally validated using new data and found to have a F value of 9.892, with a significant p-value of 0.019. A significant F value indicates that the regression model provides a better fit than a model without independent variables or simply implies that the descriptors are indeed predicting the dependent variable (Sureiman & Mangera, 2020). Additionally, the 2D-QSAR model developed had an R-squared value of 0.978 and an adjusted R-squared value of 0.8768 indicating good predictive power. Adjusted R-squared is a better version of R-squared that adjusts for the number of independent variables in a regression model and measures the goodness of fit of a regression model. Hence, a higher adjusted R-squared above 0.85 (85%) implies that model is a good fit and can predict the dependent variable effectively (Chicco et al., 2021).

Durbin Watson (DW) statistic detects autocorrelation in the residuals from a regression model. The DW statistic ranges from 0 to 4, where a value of 2.0 indicates zero autocorrelation while values below 2.0 and above 2.0 indicate positive autocorrelation and negative autocorrelation, respectively (Rajabi & Shafiei, 2019). In this study, the 2D-QSAR model had a DW value of 2.19 that was non-significant which strongly indicated zero serial correlation. When two or more independent variables are highly correlated, then multicollinearity can occur and under such scenarios, the highly correlated predictor variables don’t provide any unique information for the regression model. Multicollinearity can cause problems when fitting and interpreting the regression model. Variance inflation factor (VIF) measures the strength of correlation between the predictor variables in a regression model (Munjala et al., 2017), and the value for VIF starts at 1 and has no upper limit. VIF values are categorized as 1, between 1 and 5 and greater than 5 indicating no correlation, moderate correlation and severe correlation between predictor variables in the regression model. The 16 PADEL descriptors used to fit the model had VIF value ranging from 3.19 to 15.73, although an indication of moderate to severe multicollinearity, in some cases high VIF values can be ignored and not relied upon during model validation.

The mean absolute error (MAE) measures the accuracy of a given model. This indicates the average absolute difference between the observed values and the predicted values. Moreover, Mean Bias Error (MBE) is the mean difference between the predicted values and the actual values which indicates the ability of the measurement process to underestimate or overestimate the value of a given parameter. MBE can either be of either positive bias indicating that the error arising from the data is overestimated or negative bias indicating underestimation of MBE. MBE should be handled with care as the positive and negative errors may cancel each other out (Chicco et al., 2021). The developed model had a low MAE and MBE of 0.1748 and 6.97 x 10^−16^ respectively indicating a better-fitting model.

In addition, Relative Absolute Error (RAE) measures the performance of a predictive model and is expressed in terms of a ratio ranging from 0 to 1. RAE indicates how the mean residual relates to the mean deviation of the target function from its average. A good model will have RAE values close to zero, and zero is the best value. (Chicco et al., 2021). Furthermore, Mean Absolute Percentage Error (MAPE), also known as Mean Absolute Percentage Deviation (MAPD) increases linearly with an increase in model error. Hence, the smaller the MAPE, the better the model will perform. Moreover, Mean Squared Error (MSE) is another important model parameter since squaring the error will give a higher weight to the outliers, which results in a smooth gradient for small errors (Vishwakarma et al., 2021). The value of MSE ranges from 0 to infinity and increases exponentially with an increase in model error. A good model will have an MSE value closer to 0 while a poor model will have a higher MSE. Besides, Root Mean Square Error (RMSE), measures the average magnitude of the errors and focuses on the deviations from the actual data values. RMSE value of zero implies a perfectly fitting model hence the lower the RMSE, the better the model and its predictions while a higher RMSE indicates a poor model where there is a large deviation from the residuals and the actual values. (Quadri et al., 2022). The developed model had RAE of 0.1698, MAPE of 0.1566, MSE of 0.0469 and RMSE of 0.2166 which are low values close to 0 and all are in line with a model with a good predictive power.

### Pharmacophore modeling results

From PharmaGist Webserver analysis, the best pharmacophore features for a ligand that can bind to PBP2a were 1 aromatic ring, 1 hydrophobic group, 1 hydrogen donor, 1 negative ion and 4 hydrogen acceptors. After searching the pharmacophore in the various chemical databases using the Pharmit server (Sunseri & Koes, 2016), a list of 242 compounds (**Supplementary_data_3.zip**) fitting the pharmacophore profile was obtained and subsequently subjected to ADME screening.

### ADME Screening

The list of ligands was then subjected to ADME screening using the SwissADME (Daina et al., 2017) online tool. Ligands were screened for water solubility, oral drug absorption, lipophilicity and other pharmacokinetics like blood-brain barrier (BBB) permeation, skin permeability, gastrointestinal (GI) absorption, inhibition of cytochrome P450 enzymes and drug-likeness. Physicochemical parameters like the number of hydrogen bond acceptors and donors, number of rotatable bonds, molar refractivity and topological polar surface area, molar refractivity and fraction of carbons in the sp3 hybridization were also screened. Only compounds with promising physicochemical parameters, water solubility, lipophilicity, pharmacokinetics and medicinal chemistry (which included screening to determine whether the ligands had any PAINS and Brenk alerts, were lead-like and synthetically available) were subjected to the developed QSAR model to predict their anti-PBP2a activity (**Table 2**). In addition, the selected compounds were drug-like and fulfilled Lipinski (Pfizer) filter, the pioneer rule-of-five (Lipinski et al., 1997), Ghose (Amgen) (Ghose et al., 1999), Veber (GSK) (Veber et al., 2002), Egan (Pharmacia) (Egan et al., 2000) and Muegge (Bayer) (Muegge et al., 2001) filters for prospective oral drugs. A total of 16 compounds were selected from the original 242 compounds after ADME screening and subjected to the developed 2D-QSAR model for the prediction of anti-PBP2A activities.

**Table 2:**
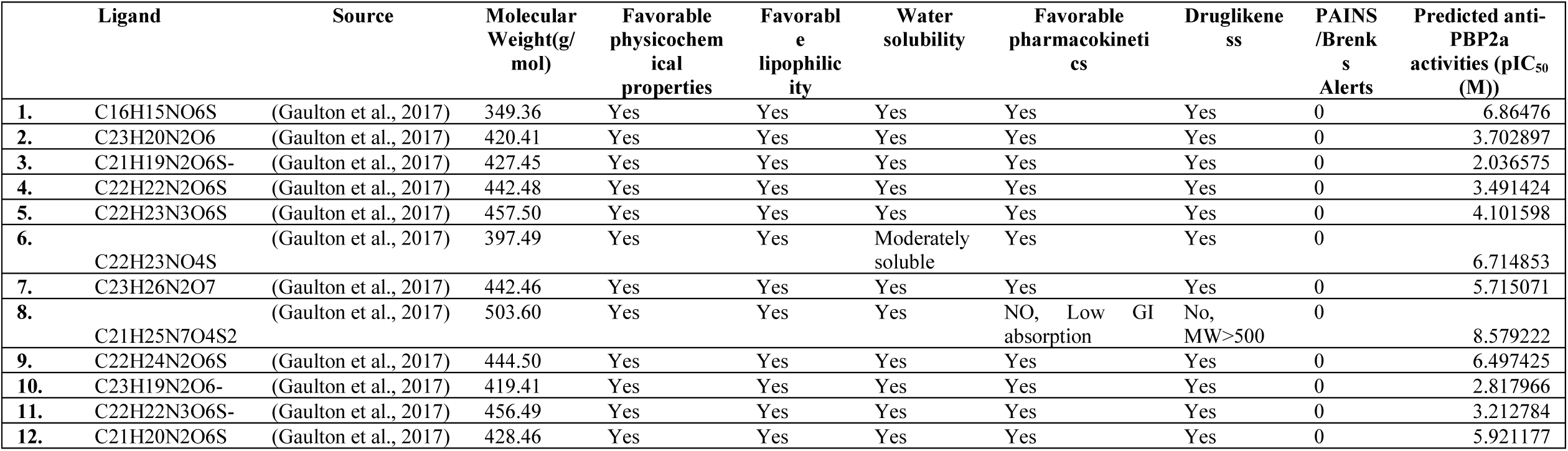
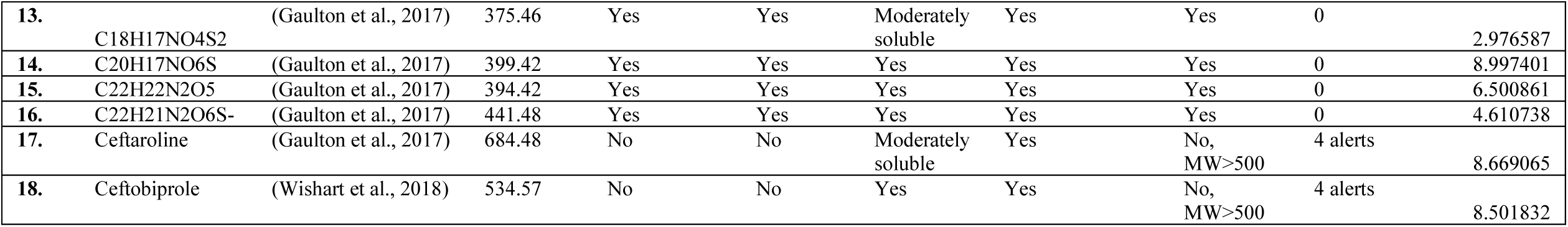
Predicted ADME properties and anti-PBP2a activities of selected ligands.

### Prediction of anti-PBP2A activity of ligands

2D descriptors for the 16 compounds (plus 2 positive controls) were generated using PaDEL followed by data cleaning through the removal of constant columns, blank entries and highly correlated columns. This new test data set was checked to ensure that it has the same number of descriptors as those used to build the model. After subjecting the new data to the model, the following anti-MRSA activities (pIC50) were predicted (**Table 2**). IC50 value measures the effectiveness of compound inhibition towards biological or biochemical utility. Occasionally, it is also converted to the pIC50 scale (−log IC50), in which higher values indicate exponentially greater potency (Hendrickx et al., 2018). pIC50 values are the experimental values which are predicted and coded in nanomolar (nM) or micromolar(µM) hence pIC50 is a new approach to examine the same data in a logarithmic manner. These IC50 values differ in a wide range of molecules in various applications of molecular modeling techniques such as QSAR. So pIC50 makes data easily comparable, understandable and eliminates the possibility of errors in data representation and reproducibility (Thakur et al., 2022). In the current study, the activity of the 16 compounds was converted into logarithmic values in molar units (pIC50 = −log (IC50*10 −9)).

Predicted ADME and anti-PBP2a activities were compared among the 16 ligands (all available in the CHEMBL database) plus two positive controls; ceftaroline and ceftobiprole. Compounds with highest predicted anti-PBP2a activities (pIC_50_) included Ligand **1 (**ChEMBL307337): **C_16_H_15_NO_6_S** (6.86476 M), ligand **6 (**[ChEMBL79476](https://www.ebi.ac.uk/chembl/explore/compound/CHEMBL307337)): **C_22_H_23_NO_4_S** (6.714853 M), ligand **8 (**ChEMBL30602): **C_21_H_25_N_7_O_4_S_2_** (8.579222 M), ligand **9 (**ChEMBL1207733): **C_22_H_24_N_2_O_6_S** (6.497425 M), ligand **14 (**ChEMBL304837): **C_20_H_17_NO_6_S** (8.997401 M), ligand **15 (**ChEMBL81062): **C_22_H_22_N_2_O_5_** (6.500861 M), **Ceftaroline** (8.669065 M) and **Ceftobiprole** (8.501832 M). The 6 ligands with high anti-PBP2a activity together with the two positive controls were subjected to molecular docking to investigate the molecular interactions between them and the PBP2a of MRSA.

### In silico anti-PBP2A activity

Six ligands with the highest anti-PBP2a activity were subjected to molecular docking and the interactions between PBP2a and the ligands with the highest anti-PBP2a activity are shown in **Table 3** and **Figure 2-4**.

**Table 3:** Binding energies and molecular interactions between ligands and PBP2a.

| Potential PBP2a inhibitor | Ligand-PBP2a Complex Binding energy [ΔG bind(kcalmol<sup>-1</sup>)] | Amino acids that interact with compounds through hydrogen bonds | Amino acids that make hydrophobic and other Interactions (van der Waals, pi-sigma bonds, alkyl, pi-alkyl, Pi-Pi T-shaped, Pi-Sigma) | Total number of interactions (Hydrogen bonds & other interactions) |
| --- | --- | --- | --- | --- |
| C<sub>16</sub>H<sub>15</sub>NO<sub>6</sub>S | -7.5 | ARG 241, HIS 293(2) | LYS 148, SER 149, ARG 151, THR 165, ASN 164, GLY166, SER 240, MET 372, PRO 258, VAL 277, GLY257, GLY 257, TRY 373, VAL 256 | 16 |
| C<sub>22</sub>H<sub>23</sub>NO<sub>4</sub>S | -8.1 | ARG 241, ASN 164, THR 165 | TRY373, MET372, GLY257, VAL256, GLY166, ASN242, ASN164, SER240, GLU150, THR165, LYS148, GLU239, SER149, HIS293, SRG151, VAL277, PRO258 | 20 |
| C<sub>21</sub>H<sub>25</sub>N<sub>7</sub>O<sub>4</sub>S<sub>2</sub> | -8.6 | HIS293, LYS148, GLU239, VALA256, SER240 | SER149, ARG151, THR165, GLU150, ARG 241, ALA168, THR216, GLY257, MET372, VAL277, PRO258 | 16 |
| C<sub>22</sub>H<sub>24</sub>N<sub>2</sub>O<sub>6</sub>S | -7.3 | THR165, GLU239 | TRY373, VAL256, ARG241, GLU150, SER149, HIS293, LYS148, ASN164, THR 238, ARG151, VAL277 | 13 |
| C<sub>20</sub>H<sub>17</sub>NO<sub>6</sub>S | -8.3 | HIS293, ARG241(2), THR165 | PRO258, TRY373, MET372, GLY257, LYS148, VAL277, VAL256, GLU150, SER149, GLU239, ARG151, GLY166, ASN164 | 16 |
| C<sub>22</sub>H<sub>22</sub>N<sub>2</sub>O<sub>5</sub> | -8.0 | ASN164, ARG241(2), THR165 | GLY257, MET372, PRO258, TYR373, VSL256, VAL277, HIS293, SER240, LYS148, GLU239, GLU150, ARG151, GLY166, ASN242 | 18 |
| Ceftaroline | -8.0 | LYS153, GLU239(2), LYS318, GLU150, THR165 | ASP320, GLU150, ARG151, SER240, HIS293, VAL277, LYS272, GLU294, ALA276, ASP275, LYS148, LEU147, LYS317, GLU239 | 20 |
| Ceftobiprole | -8.8 | GLU170, VAL256, MET372, VAL277 | PRO258, GLY257, SER240, ASP275, ALA276, LYS273, GLU294, ASP295, SER149, LYS148, HIS293, ARG151, GLU150 | 17 |

All the 6 ligands displayed favorable binding affinities with PBP2a receptor of around −8.0 kcalmol^−^ which was similar to that of positive controls of ceftaroline (−8.0 kcalmol^−1^) and ceftobiprole (−8.8 kcalmol^−1^). The ligands displayed favorable hydrogen bonding besides other interactions revealing favorable docking scores. Based on the binding affinities, two ligands **8**: **C_21_H_25_N_7_O_4_S_2_** (8.579222 M) and **14: C_20_H_17_NO_6_S** (8.997401 M) displayed the highest docking scores of -**8.6 kcalmol^−1^**and **-8.3 kcalmol^−1^** respectively that were not significantly different from the positive controls’ docking scores. 2D structures of ligands **8**, **14** and ceftobiprole are displayed in Figure **1**. The interactions of PBP2a with ligands **8** and **14** suggest that the ligands have potential anti-PBP2a activities (**Figure 2 & 3**).

**Figure 1:**
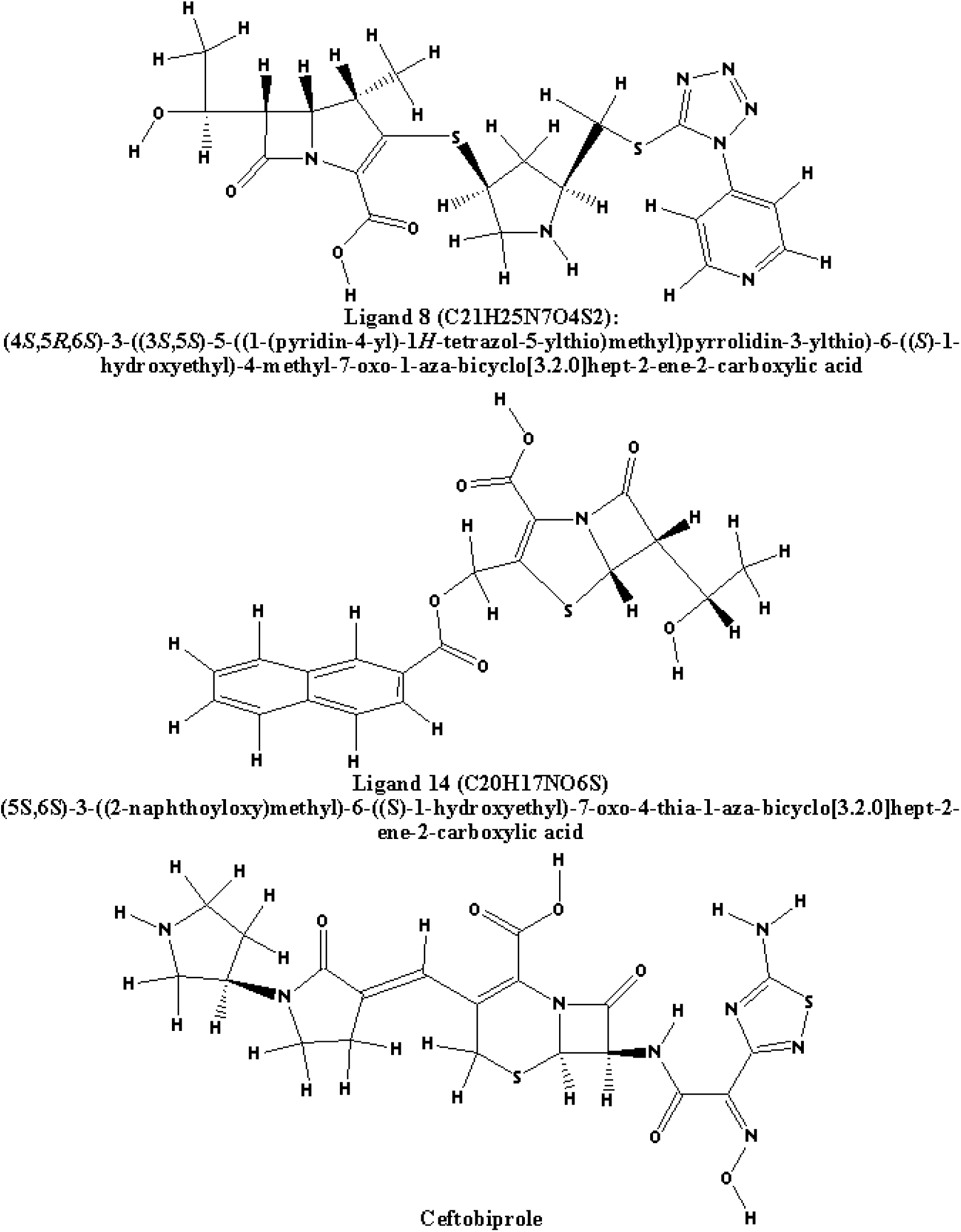
2D structure of Ligand **8**, **14** and **Ceftobiprole**.

**Figure 2:**
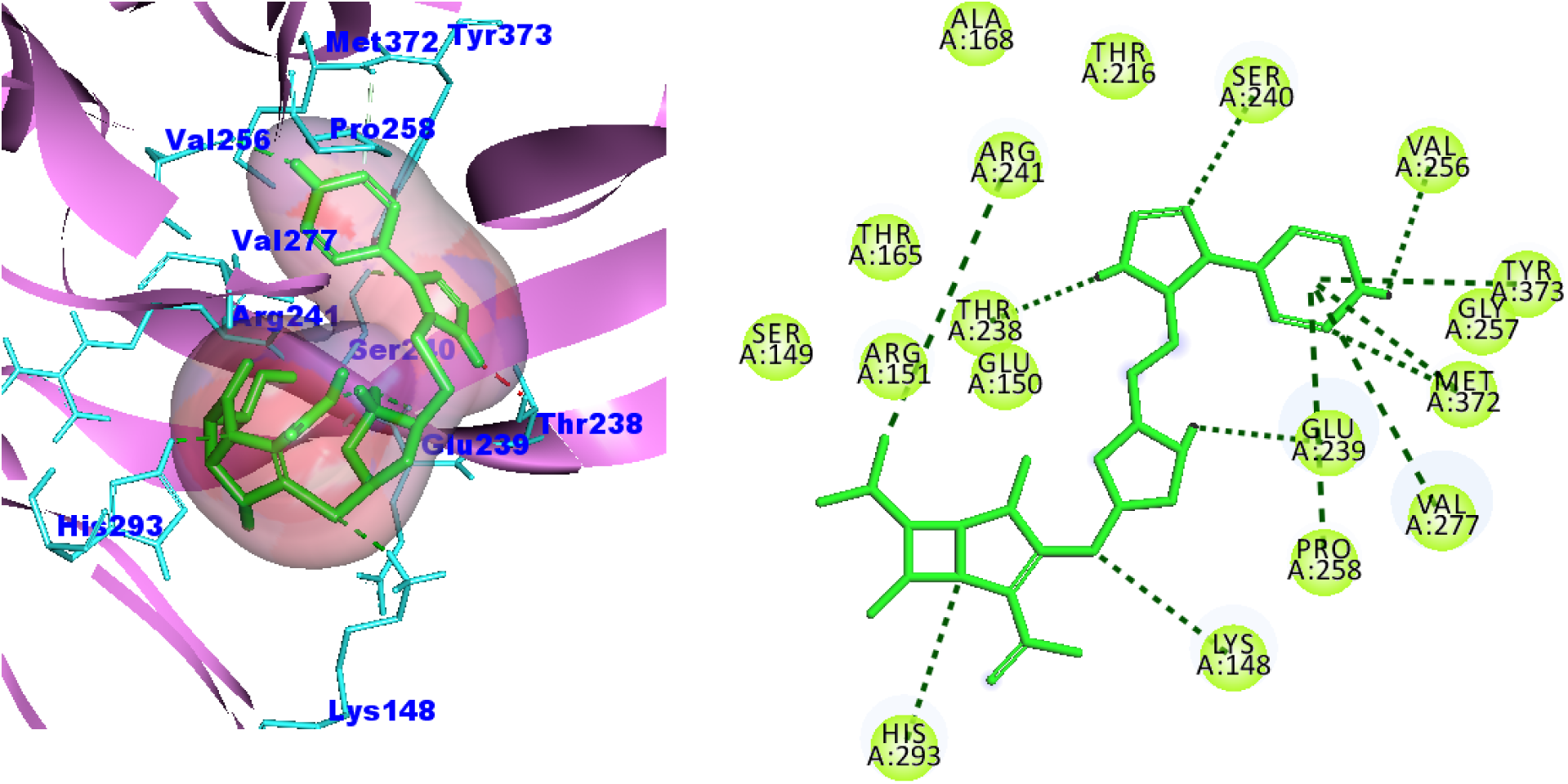
Interactions of Ligand **8** with PBP2a: Interactions include: Van der Waals forces observed in SER149, THR165, ARG151, GLU150, ALA168, THR216 and GLY257, conventional hydrogen bond seen in HIS293, GLU239, LYS148, VAL256, SER240, Pi-Pi T-shaped, Alkyl and Pi-Alkyl in TRY373, ARG241, VAL277 and PRO258Binding energy= −8.6 kcalmol^−1^.

**Figure 3:**
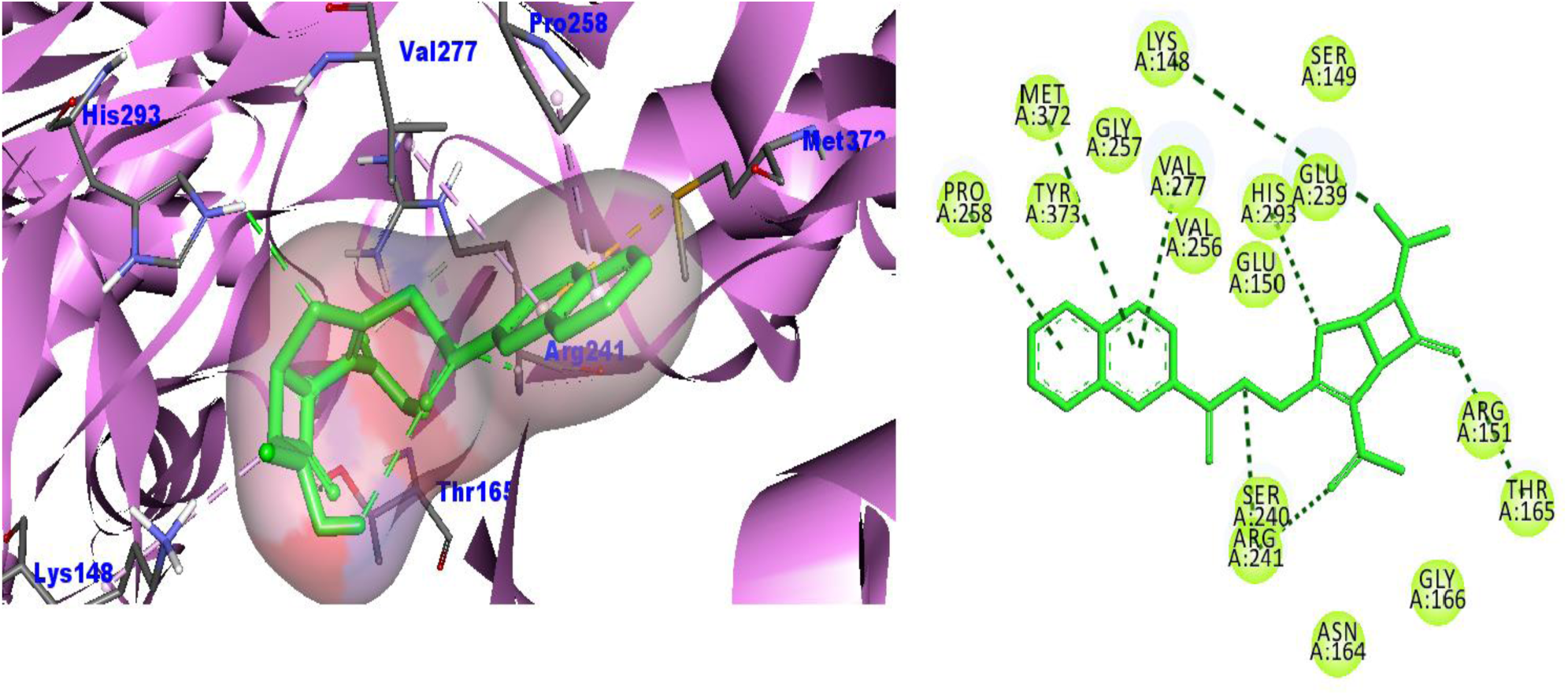
Interactions of Ligand **14** with PBP2a of MRSA. Interactions include: Van der Waals forces observed in TRY373, GLY257, VAL256, GLU150, GLU239, SER149, SER 240, GLY166 and ASN164, conventional hydrogen bond seen in ARG241, THR165, HIS293, pi-Sulphur bonds seen in MET372, alkyl and pi-Alkyl interactions observed in PRO258, VAL277 and LYS 148. Binding energy= −8.0 kcalmol^−1^.

The pharmacophore maps and QSAR models can be used to come up with novel and more active analogs. The pharmacophore models which define noncovalent and covalent interactions with receptors can be generated based on different properties such as hydrogen bond donor, hydrophobic group, hydrogen bond acceptor, positive and negative ionic features from analysis of several compounds which bind to the same receptor. Some pharmacophore features are favorable while others are unfavorable that may be unnecessary for ligand-receptor interactions hence a model with favorable features is desired (Tawari et al., 2008). QSAR models have successfully been used in the identification of novel compounds with various inhibitory activities against the same targets. Further understanding of the inhibitory activity can be gained by envisaging QSAR models in the context of ligands in a series with varying activities on the same receptor (Veerasamy et al., 2011). The established QSAR model was externally validated using 16 test set molecules, which gave an excellent value of 0.879 (>0.5), found to have a F value of 9.892, with a significant p-value of 0.019, R^2^ of 0.978, Multiple Adjusted R^2^ of 0. 8768, Durbin Watson (DW) statistic of 2.19 with a p-value of 0.864, Mean absolute error (MAE) of 0.1748, Mean Bias Error (MBE) of 6.97 x 10^−16^, Relative Absolute Error (RAE) of 0.1698, Mean Absolute Percentage Error (MAPE) of 0.1566, Mean Squared Error (MSE) of 0.0469 and Relative Mean Squared Error (RMSE) of 0.2166. External validation, indicated that the QSAR model possessed a high accommodating capacity, it can be reliable and can be used to predict the anti-PBP2a activities of molecules.

The top pharmacophore model generated from the features of 7 molecules with good binding affinity for PBP2a was found to be associated with seven-point hypotheses which were: 1 aromatic ring, 1 hydrophobic group, 1 hydrogen donor, 1 negative ion and 4 hydrogen acceptors. Such a pharmacophore model was used to search for other molecules across different databases using the Pharmit server. A total of 242 molecules were obtained from the various databases and subjected to ADME screening. Eventually 16 molecules that had favorable ADME properties were selected from the 242 compounds. Similarly, molecules that specifically target PBP2a of MRSA were obtained from the BindinBD data, their descriptors were calculated and 2D-QSAR model was generated in R. Molecules searched using the pharmacophore model were subjected to the QSAR model to predict their anti-PBP2a activity then the most active molecule were subjected to molecular docking to study the molecular interactions between them and PBP2a.

A molecular docking study was carried out on the 6 compounds predicted to have high anti-PBP2a activity based on the 2D-QSAR model developed. Results indicated that ligands **8** and **14** interacted with the PBP2a receptor better compared to the other compounds. Indeed, the two compounds had a binding score of −8.6 and −8.3 kcalmol^−1^ respectively when docked to the active site of the PBP2a of MRSA. Such docking score was similar to the docking score observed when ceftobiprole and ceftaroline were docked to the active site of the PBP2a of MRSA. Such high docking scores were attributed to the hydrogen bonding, hydrophobic and other interactions such as van der Waals, pi-sigma bonds, alkyl, pi-alkyl, Pi-Anion, Pi-Pi T-shaped, Pi-Sigma interactions. More interactions mean better docking score, in total, both ligands **8** and **14** were able to bind to PBP2A receptor through 16 different interactions as shown in **Table 3**. More hydrogen bonds in the ligand-receptor complex are preferred and enhance ligand-receptor interactions resulting in better docking scores (Chen et al., 2016). The most common amino acids of PBP2a which formed hydrogen bonding with ligand **8:** C_21_H_25_N_7_O_4_S_2_ (ChEMBL30602) and ligand **14**: C_20_H_17_NO_6_S (ChEMBL304837) were HIS293, LYS148, GLU239, VAL256, SER240, ARG241 and THR165 as shown in Figures 2-4.

**Figure 4:**
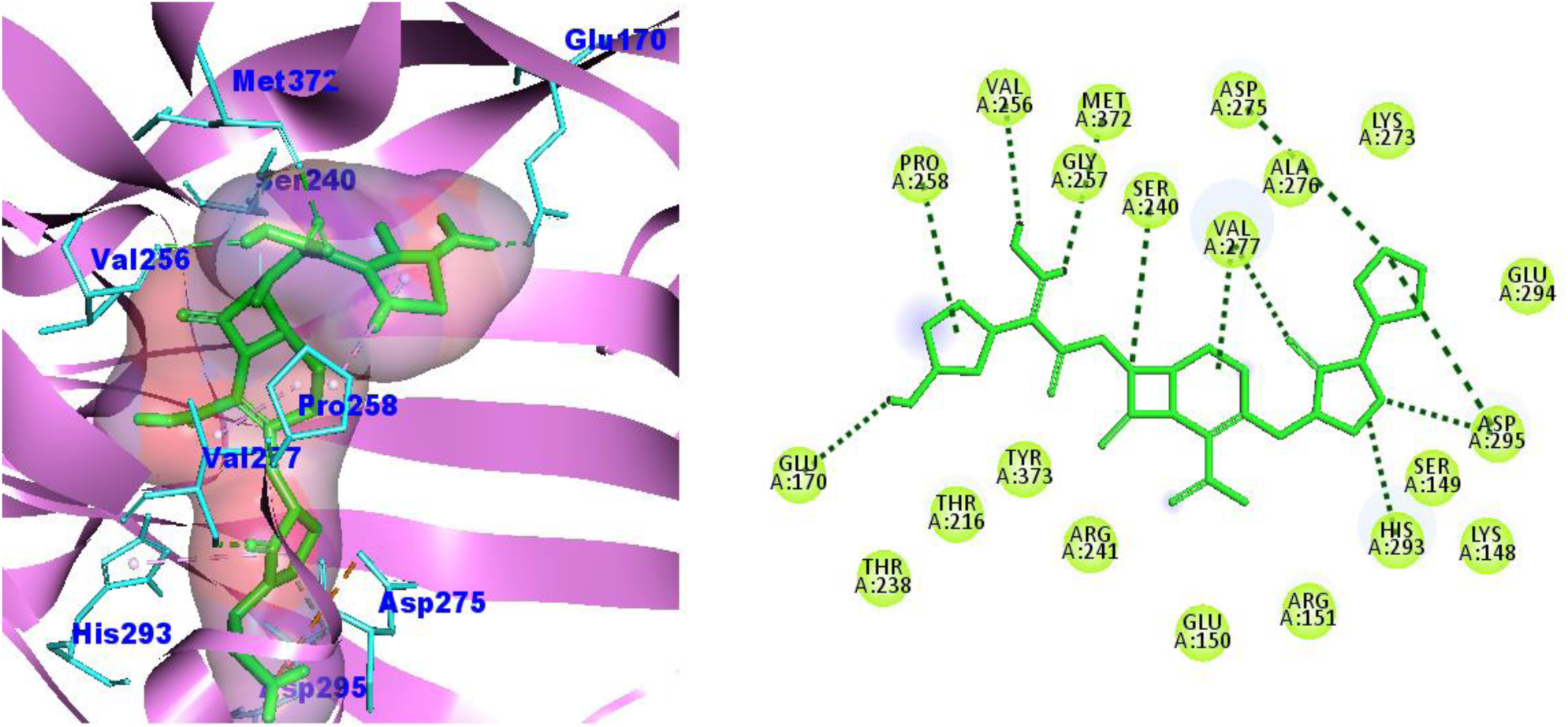
Interactions of Ceftobiprole with PBP2a of MRSA. Interactions include: Van der Waals forces observed in THR238, THR216, TYR373, ARG241, GLU150, ARG151, LYS148, SER149, GLU294, LYS273, ALA276, SER240 and GLY257, conventional hydrogen bond seen in GLU170, VAL256, MET372, SER240 and VAL277, attractive charges, alkyl and pi-alkyl bonds seen in PRO 258, ASP275, ASP295, VAL277 and HIS 293; Binding energy= −8.8 kcalmol^−1^.

Integration of pharmacophore modeling and statistically significant predictable QSAR provides information about the structure and the relationship between the structure and activities of the molecules under study (Kirubakaran et al., 2012). This will provide in-depth information related to *in silico* structure editing and modifications of some structural side chains which can aid design of novel molecules with better activities before synthesis (Silverman & Holladay, 2014). Molecular docking studies also support by confirming the results observed from pharmacophore maps and QSAR models by showing the binding interactions and orientation of these potential inhibitors are compared with the native ligands of PBP2a such as ceftobiprole and ceftaroline. The results of this study indicate that the 6 molecules identified possess high anti-PBP2a activity and also fit and bind the binding pocket of the PBP2a receptor of MRSA.

### MD simulation analysis

From molecular docking and QSAR prediction results, ligands 8 and 14 were shown to possess high anti-PBP2a activity hence both ligands together with the positive control ceftobiprole were subjected to MD simulation over 100 ns. The RMSD of the protein was calculated over a 100 ns when bound to the 3 compounds (ligand 8, 14 and ceftobiprole). Figure 5A shows the RMSD plot of the protein Cα atoms, while Figure 5B shows the RMSD of the ligands when they are bound to the protein. The RMSD of the protein was stable for most of the complexes. The RMSD of the protein increased from its initial state from 0.2 nm to 0.55 nm. Moreover, in ceftobiprole, there are several peaks that reach around 0.6 nm, indicating periods of higher deviation from the reference structure around 70 ns. Similarly, ligand 8 showed relatively consistent RMSD values around 0.4 to 0.5 nm with fewer large deviations at 70ns. Ligand 14 has the lowest RMSD values overall, mostly staying between 0.2 and 0.4 nm, indicating the least deviation and potentially the most stable interaction with the protein. Overall, the outcome suggested the ligand-protein complex with ligand 14 appears to be the most stable, with the lowest and most consistent RMSD values. The RMSD of the ligand is shown in Figure 5B. Here, ceftobiprole and ligand14 showed stable conformation. Ligand 8 showed a high deviation after 50 ns of the MD run; therefore, it was excluded from further analysis. The RMSD of the other two ligands, ligand 14 and ceftobiprole, was calculated and is given in Figure 5C and the RMSD for both ligands were found to be stable as it was below 0.3 nm as shown in the figure. Both ligands have an initial RMSD of 0.5 nm, the RMSD begins around 0.5 nm and increases over time, stabilizing around 1.5 nm with fluctuations which suggests that ceftobiprole undergoes more significant conformational changes compared to ligand 14. Similarly, the ligand 14 exhibits stable conformation up until 30 ns, at which point it deviates more to 1 nm, drops to 0.5 nm, and then exhibits a substantial deviation at the simulation’s end. The root mean square fluctuation (RMSF) of the protein and ligand was calculated to quantify the variations of each residue in the bound state. The RMSF of the protein when in a bound state is shown in Figure 5D. The majority of the residues of the protein showed RMSF of < 0.6 nm. Few fluctuations (2–3 peaks) were observed of > 0.6 nm during the 100 ns of the simulation which indicated stable interactions between the protein and ligands under study.

**Figure 5.**
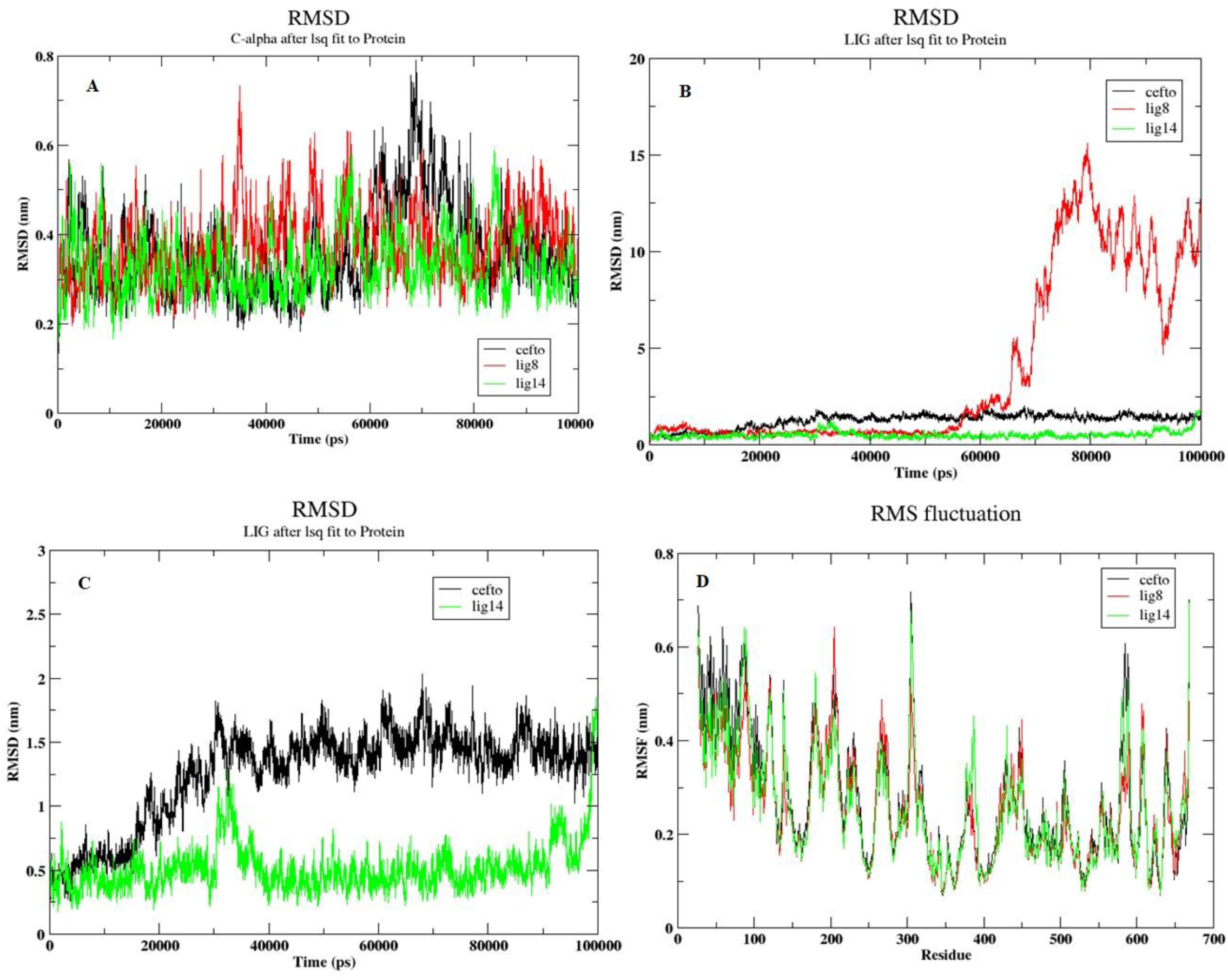
**A**: RMSD of the protein C-alpha atoms, **B**: RMSD of the ligands when bound with the protein, **C**: RMSD of the ligands when bound with the protein, **D**: RMSF of the protein residues when bound with the ligands, cefto=ceftobiprole, lig8=ligand 8, lig14=ligand 14

The protein’s SASA averaged for all residues during the 100 ns simulation shown in Figure 6 **A**. The initial SASA of the ligand 14 was 330 nm^2^, and for the ceftobiprole, it was 335 nm^2^. The SASA value fluctuated between 330 nm^2^ and 350 nm2 throughout the simulation. It appears that ligand 14 has a more stable interaction with the protein. While ceftobiprole, shows more variability in solvent exposure, which may indicate more dynamic behavior or less stable binding. The radius of gyration (Rg) is shown in Figure 6B, where the compactness of the protein is determined. The ligands ceftobiprole and ligand 14 fluctuate around a similar range, indicating that both molecules have similar levels of compactness or extension during the simulation. The Rg for both molecules appear to fluctuate between approximately 3.6 nm and 3.8 nm, with occasional peaks. This indicates that both ceftobiprole and ligand 14 exhibit similar levels of structural compactness with continuous fluctuations throughout the simulation period. Hydrogen bonds were calculated between the protein and ligand molecules and were identified. The ligand 14 shown in Figure 6C forms a 2–3 hydrogen bond. There are occasional spikes where the number of hydrogen bonds reaches up to 4 or 5, but these are relatively rare. The number of hydrogen bonds in ceftobiprole (Figure 6D) also fluctuates, but with a different pattern compared to ligand 14. Initially, there are higher spikes, with the number of hydrogen bonds reaching up to 6 within the first 20,000 ps. After this initial phase, the number of hydrogen bonds stabilizes more around 1 or 2, with occasional drops to 0 or increases up to 4. The plots reveal that ligand 14 maintains a steady, though fluctuating, pattern of hydrogen bonding, while ceftobiprole exhibits an initial high activity followed by a more stable period with fewer hydrogen bonds.

**Figure 6A:**
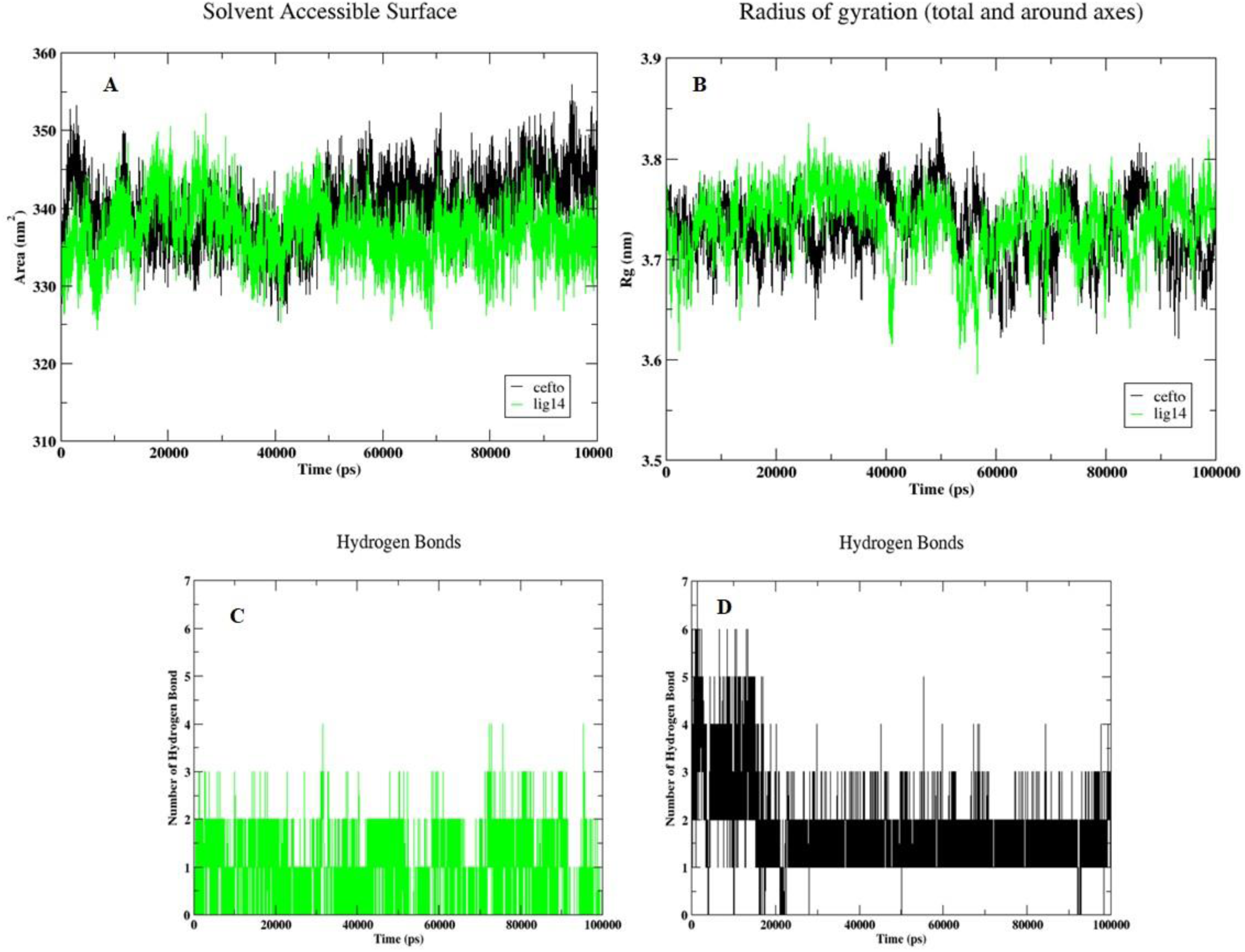
Solvent accessible surface area (SASA) of every residue of protein, **B**: Radius of gyration of protein in bound state with the ligand for 100 ns of simulation, **C & D**: Hydrogen bonds formed between protein and ligand during 100 ns of simulation for ligand 14 and ceftobiprole respectively. cefto=ceftobiprole, lig14=ligand 14

The principal component analysis (PCA) scatter plot for ceftobiprole exhibits a wide dispersion of states, indicating a diverse range of conformations explored during the 100 ns simulation. The PCA is given in the Figure 7A where the distribution of the black dots for ceftobiprole seems to be more dispersed, covering a larger area in the space, suggesting a broader range of conformational states. The green dots for ligand 14 are also widely spread but tend to cluster more around the center, indicating that ligand 14 might have more constrained conformational variability compared to ceftobiprole. Overall, it suggests that ligand 14 maintains more consistent conformations and potentially more stable interactions with the protein. While ceftobiprole indicates greater conformational flexibility or instability in its binding interactions with the protein. In **Figure 7B & C**, the plot shows a central region in dark blue, indicating a low free energy area. The Free energy landscape (FEL) plots for ligand 14 and ceftobiprole showed that both molecules have stable conformations represented by the dark blue central regions. Ligand 14 has a more compact, stable region with smoother transitions to higher energy states, while ceftobiprole shows a broader range of stable conformations and more pronounced energy barriers. The binding free energy of the complexes was calculated using the MM/GBSA technique for the entire 100 ns trajectories. It was observed that ceftobiprole-PBP2A complex had the lowest binding free energy compared to ligand 14-PBP2A complex. Here, ceftobiprole showed a total binding energy of −16.90 kcal/mol, while ligand 14 had a total binding energy of −12.38 kcal/mol. The binding affinities over the entire MD run for both ligands strongly indicate that they have robust affinity for PBP2a.

**Figure 7A:**
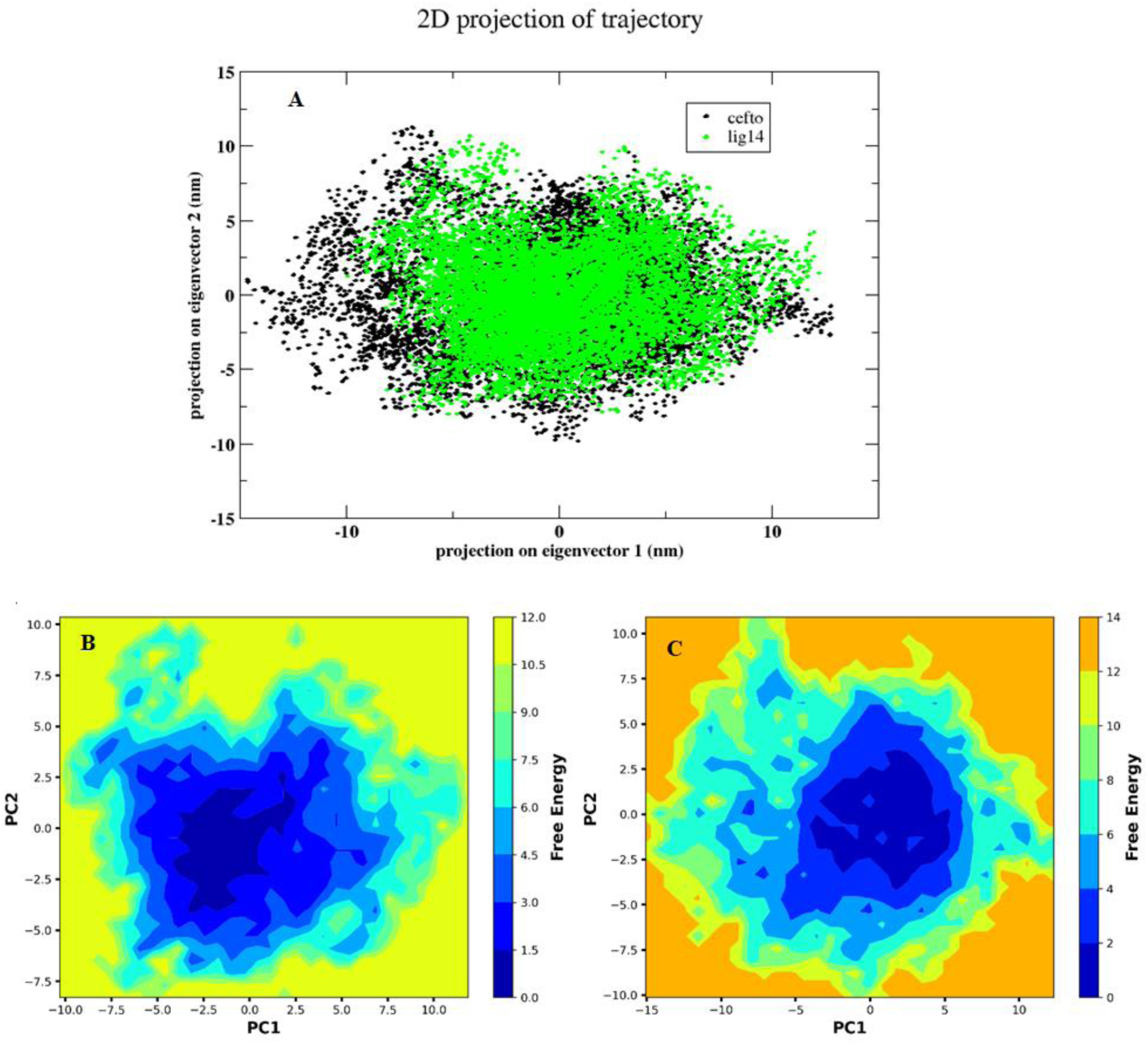
Principal component analysis (PCA), green dot shows the ligand 14 and black dots shows the ligand ceftobiprole, Free energy landscape (FEL) (**B**)ligand 14 (**C**)ceftobiprole

Overall, in this study, it was observed that ligand 8 exhibited a higher RMSD compared to ligand 14 and ceftobiprole. Due to this high RMSD, ligand 8 was excluded from further analysis. Ligand 14 and ceftobiprole showed a low RMSD hence are able to form a stable complex bound to PBP2A. In the hydrogen bond analysis, it was observed that ceftobiprole formed more hydrogen bonds throughout the simulation than ligand 14. This indicates stronger or more frequent interactions between ceftobiprole and PBP2a receptor. Similarly, in the PCA, both ceftobiprole and ligand 14 displayed a range of structural variability suggesting both ligands explore a wider range of conformational states during the simulation. Additionally, both ligands were found to have robust binding affinity for PBP2A based on the free energy calculations. Inclusively, these analyses highlight that ligand 14 and ceftobiprole form stable interactions and have strong binding affinity for PBP2A.

## Conclusion and Recommendations

The current study involved, ligand-based pharmacophore modeling, atom-based 2D-QSAR model and ADME screening. The outcome was the identification of new molecules with favorable pharmacophore features and ADME properties with the potential to bind to PBP2a of MRSA as well as the development of a satisfactory, predictive and significant QSAR model for predicting anti-MRSA activities of new compounds. High affinity to the PBP2a receptor was also confirmed through molecular docking. Of particular interest were two compounds that showed high predicted anti-PBP2a activities based on the developed QSAR model and molecular docking study. The compounds were ligand **8**: ((4S,5R,6S) −3- ((3S,5S) −5- ((1-(pyridin-4-yl) - 1H-tetrazol-5-ylthio) methyl) pyrrolidin-3-ylthio) −6- ((S) −1-hydroxyethyl) −4-methyl-7-oxo-1-aza-bicyclo [3.2.0] hept-2-ene-2-carboxylic acid) and ligand **14:** ((5S,6S)-3-((2-naphthoyloxy) methyl) −6-((S)-1-hydroxyethyl) −7-oxo-4-thia-1-aza-bicyclo [3.2.0] hept-2-ene-2-carboxylic acid). From MD simulation analysis, it was observed that ligand **14** has favorable interactions with PBP2A based on RMSA, RMSF, Rg, SASA, PCA and binding energies and its interactions with PBP2A is similar to the interactions of positive control drug ceftobiprole with PBP2A. Hence, ligand **14**: C_20_H_17_NO_6_S (ChEMBL304837) may become potential drug candidate against MRSA which has developed a lot of resistance to current antibiotics. However, further *in vitro* and *in vivo* studies of the compound is required to validate these *in silico* findings

## Data Availability

Data used in this research work is available from the corresponding author upon request.

## Authors’ contributions

The following authors contributed to this study: conceptualization, F.M.M and G.W.G; methodology, F.M.M, G.W.G and P.W.A; data analysis and MD simulation, D.B.K, F.M.M and P.N.N; writing-original draft preparation, G.W.G and F.M.M; writing, review and editing, P.W.A, G.W.G, J.M.K, D.B.K, F.M.M and P.N.N. All authors have read and agreed to the published version of the manuscript.

